# The heterogeneity of age-related muscle atrophy of lower limb muscles

**DOI:** 10.64898/2026.09.15.751873

**Authors:** Christopher D Connelly, Jonathan P Folland, Mathew Piasecki, Jakob Škarabot

## Abstract

Age-related skeletal muscle atrophy differs between upper and lower body muscles and may also differ between lower limb muscles, but these muscle-specific patterns remain poorly characterised. We acquired axial 3T magnetic resonance images from sex-balanced samples of thirty young (24±4 years) and thirty older adults (73±4 years). Muscle volume of seventeen lower limb muscles/compartments was quantified using manual segmentation and expressed as absolute volume, relative to body mass, and as a percentage of total muscle volume. Absolute muscle volume was lower in older adults in 13 out of 17 muscles compared to young, with substantial heterogeneity in the effect sizes (*d*=0.01-3.36). Age group × muscle interaction contrasts revealed muscle-specific age-related differences in volume. Three of the quadriceps muscles demonstrated greater age-related atrophy compared to several other muscles (e.g. vastus lateralis differing from 14 of the other 16 muscles; *d*≥1.54). Conversely, the anterior and deep posterior compartments of the shank and biceps femoris short head showed significantly smaller age-related atrophy than several other muscles. These interaction effects were exacerbated when expressed in relative terms. Consequently, three of four shank compartments, gluteus maximus and adductor magnus represented a greater proportion of total muscle volume in older adults, whereas the vastus lateralis, vastus intermedius, rectus femoris, semitendinosus, gracilis, and iliopsoas contributed a smaller proportion in older adults. These findings highlight the variability in age-related leg muscle atrophy in a large mixed sex cohort, resulting in a redistribution of muscle volume in the lower limb.

## Introduction

Ageing is associated with a decline in skeletal muscle mass, contributing to the loss of physical function and mobility [1]. Although age-related muscle atrophy is well established, recent evidence suggests that the loss of muscle mass does not occur uniformly across muscles [2,3]. Such heterogeneity may have important functional consequences, as a redistribution of muscle mass could alter force production, gait mechanics, and balance control in older adults. Furthermore, understanding the extent of age-related atrophy, and its heterogeneity across muscles may be useful for prescribing exercise or other intervention strategies to preserve muscle mass and hence physical function.

It is apparent that the extent of muscle atrophy is non-uniform, with lower limb musculature exhibiting greater age-related declines than upper limb musculature [4]. Within the lower limb there also appears to be variability between muscles with different studies that used ‘gold-standard’ magnetic resonance imaging (MRI) reporting ∼30% smaller quadriceps [5], but ∼17.5% smaller triceps surae [6] in older compared to young adults. However, such between-study comparisons may be confounded by between-study variability (e.g. in the specific age groups studied, eligibility and physical activity criteria) limiting generalisability. Recently, Fuchs et al. [7] explored the variability in age-related differences in the size of a wide range of lower limb muscles and reported that, compared to young adults, older adults had ∼24% lower thigh muscle volume and ∼15% lower pelvic muscle volume, while lower leg muscle volume did not differ between groups. However, this sample was limited to a small number of males (n = 10 in each age group), again limiting generalisability and statistical power for assessing any differential effects of age on individual muscles.

The way in which muscle size is expressed can also influence the conclusions drawn about the heterogeneity of the age-related muscle loss. Whilst absolute volume captures the total extent of muscle tissue, it is strongly influenced by body size/mass and therefore limits interpretation across individuals. Normalising muscle volume to body mass accounts for this variability in body size and thus could be a more sensitive measure of age-related changes. Indeed, relative skeletal muscle size has been reported to decline from the third decade, whereas declines in absolute size appear to be evident only at the end of the fifth decade [4]. Relative muscle size may also be a more functionally relevant index given that the lower body muscles are required to move body mass for locomotion and mobility. Finally, expressing muscle volume of each muscle/muscle group as a proportion of total muscle volume provides a measure of muscle distribution, independent of overall size [8]. For example, if the extent of age-related muscle atrophy is muscle specific [2], there may also be a redistribution of muscle mass with age. Therefore, expressing muscle volume relative to total leg muscle volume, could provide complementary information about the distribution of muscle mass with age.

Therefore, the aim of this study was to compare the muscle volume of seventeen individual lower body muscles/compartments between young and older adults and to assess whether the extent of age-related differences vary across muscles. It was hypothesised that larger proximal muscles, such as the individual quadriceps muscles, would demonstrate a greater age-related difference compared to individual distal (shank) muscles/compartments [5,7].

## Methods

### Participants

A total of 60 participant volunteered to participate in the study, consisting of 30 young (15 females; mean ± SD, age: 24 ± 4 years, height: 1.72 ± 0.10 m, mass: 67 ± 11 kg, body mass index: 22.5 ± 2.0 kg/m^2^) and 30 older adults (15 females, age: 73 ± 4 years, height: 1.66 ± 0.09 m, mass: 73 ± 11 kg, body mass index: 26.2 ± 2.7 kg/m^2^). Older adults were shorter (p = 0.0357), with a greater body mass (p = 0.0469), and body mass index (p < 0.0001) compared to young adults. All participants gave written informed consent before commencing study procedures. For eligibility, young participants were required to be aged 18-35 years, and older participants were required to be aged 65-90 years. Ineligibility criteria also included those involved in systematic training or competition, including endurance or resistance training and team sports. Additionally, participants were required to be free of neurological, cardiovascular or respiratory disorders, or lower limb musculoskeletal injury. Ethics approval was obtained from Loughborough University Ethics Committee (2021-5361-4724) and study testing was conducted in accordance with the *Declaration of Helsinki*.

Physical activity levels were estimated using the International Physical Activity Questionnaire [9]. There was no difference in estimated physical activity levels between young and older adults (young: 3034 ± 1710 vs. older: 3948 ± 2245 MET.min/week; p = 0.082). Older participants frailty phenotype status was assessed as per the guidelines of Fried et al. [10], this consisted of five criteria as follows: 1) shrinking: assessed by question “have you lost >10 lbs unintentionally in prior year”, 2) weakness: assessed by grip strength (Jamar Hydraulic Hand Dynamometer, set to position two) in lowest quartile by sex and body mass index, 3) exhaustion: assessed by answering ‘yes >3-4 days per week’ to either statement ‘I felt that everything I did was an effort’ or ‘I could not get going’, 4) slowness: assessed via 4 m walk test with gait speed in the slowest quartile by sex and height, and 5) physical activity, assessed with IPAQ score indicating <383 kcals expended per week in males and <270 kcals per week in females. If older participants met ≥3 of the criteria they were classified as frail, pre-frail if they met 1-2 and robust if they did not meet any of the criteria. Accordingly, 26 older adults were classified as robust, 4 were pre-frail and none were frail.

### Study protocols

All participants were informed to arrive for MRI scanning well rested and without having performed vigorous exercise on the day of scanning or the preceding day. A 3T MRI scanner (Discovery MR750w; GE Healthcare, Chicago, IL, USA) was used to acquire T1 axial images from the twelfth thoracic vertebrae to the calcaneus of both legs. Participants lay supine in the scanner in a full body coil with both the hips and knees fully extended, the ankles at 90^°^ and arms folded over the chest. Images were captured in five partially overlapping blocks (time of repetition 600 ms, time of echo 8 ms, field of view 450 x 450 mm, image matrix 320 x 320, pixel size 1.4 x 1.4 mm, slice thickness 5 mm, interslice gap 5 mm). On the right leg of all participants, oil filled capsules were affixed at equidistant intervals to aid the alignment of blocks for analysis.

### Magnetic resonance imaging analysis

Analysis of MR images was performed for the dominant leg/side only using an open access DICOM image-analysis software (HOROS, version 2.2.0, www.thehorosproject.org). The following seventeen muscles/muscle compartments were analysed: anterior compartment of the shank (tibialis anterior, extensor hallucis longus, extensor digitorum longus), lateral compartment of the shank (peroneous longus, peroneous brevis), deep posterior compartment of the shank (tibialis posterior, flexor hallucis longus, flexor digitorum longus, popliteus), superficial posterior compartment of the shank (soleus, medial gastrocnemius, lateral gastrocnemius, plantaris), vastus lateralis, vastus intermedius, vastus medialis, rectus femoris, semitendinosus, semimembranosus, biceps femoris long head, biceps femoris short head, sartorius, gracilis, adductor magnus, iliopsoas and gluteus maximus. Each muscle/compartment was manually segmented creating a region of interest (anatomical cross-sectional area; ACSA) on the axial slice, with this process repeated every 20 mm along the length of each muscle (Figure 1). All muscle regions of interest were overlaid onto the images to ensure there was no overlap between regions. To account for spacing between slices, a cubic spline function was fitted to the cross-sectional area data, generating 1000 interpolated values along the muscle length (R2021b; Mathworks Inc., Natick, MA, USA). Muscle volume was determined by integrating the cubic spline fitted to the ACSA-muscle length relationship; expressed in absolute terms (cm^3^), relative to body mass (cm^3^/kg), and as a percentage of total muscle volume (%).

**Figure 1.**
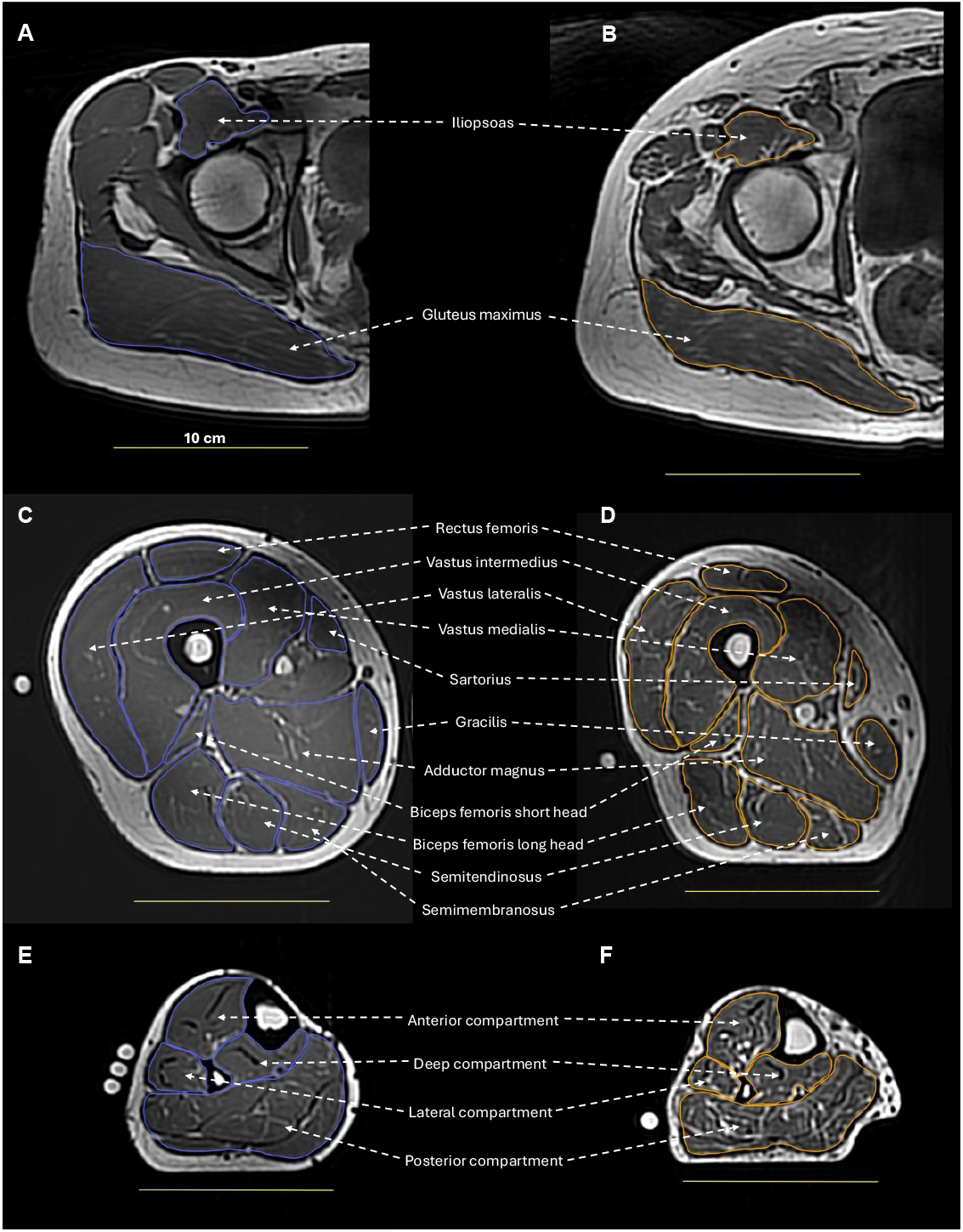
Manual segmentation of seventeen lower limb muscles/muscle compartments in a young (A, C, E) and older adult (B, D, F) from axial MR images. Images A, C, E are from a young male (age: 24 years, body mass: 76 kg, height: 1.80 cm) and B, D, F are from an older male (age: 77 years, body mass: 87 kg, height: 1.76 cm). Yellow line in each panel represents 10 cm.

### Statistical analysis

For all statistical analyses R was used (R studio, v 1.4.1106, R Foundation for Statistical Computing, Vienna, Austria). To compare descriptive participant information between young and older adults including height, body mass, body mass index, and IPAQ scores, linear models were used. To assess age-related differences in absolute and relative muscle volume, linear mixed effects models were used which included the fixed effects of age group and muscle, sex as a covariate, and participant identifier as a random intercept. However, for analysis of percentage contribution to total lower limb muscle volume, the participant random effect was estimated as zero, resulting in a singular fit. Therefore, a linear model (i.e. without a participant random effect) was used. Linear models were also constructed for comparing total muscle volume in absolute and relative terms with only the fixed effect of age group and covariate of sex. Comparison of sex differences was not directly assessed as a research question in the present study but was included as a covariate due to preliminary evidence suggesting potential differences in the trajectory of the age-related muscle atrophy between sexes [11]. Model assumptions of normality and linearity of residuals were tested using quintile-quintile plots and histograms. A square-root transformation was applied to the following variables due to non-normal distribution of residuals: absolute muscle volume, relative muscle volume, and percentage contribution to total lower limb muscle volume.

Type II Wald chi-square ANOVA using the *car* package (ver. 3.1.3) was conducted to assess model significance of the factors of age group and muscle and their interaction. To compare age-related differences in each muscle, pairwise comparisons of estimated marginal means (*emmeans* package; ver. 1.11.1, [12]) were used with Bonferroni correction for multiple comparisons. Interaction contrast testing of estimated marginal means was also performed to assess if age group differences varied across muscles, also with Bonferroni correction. The effect size of all comparisons was estimated using Cohen’s *d*, which was derived by dividing the differences in estimated marginal means by the residual standard deviation from the linear mixed effects model. Data was regarded as statistically significant with a p-value < 0.05.

## Results

Overall, older adults had a ∼21% lower total muscle volume on average compared to young adults in absolute terms (χ^2^(1) = 30.8, p < 0.0001) and ∼27% lower in relative terms, i.e. when normalised to body mass (χ^2^(1) = 132.1, p < 0.0001; Table 1).

**Table 1.** Lower limb muscle volumes in young and older adults, expressed in absolute terms, relative to body mass, and as a proportion of total muscle volume.

| Muscle/<br>muscle compartment | Absolute muscle volume (cm<sup>3</sup>) |  |  |  | Muscle volume relative to body mass (cm<sup>3</sup>/kg) |  |  |  | Percent of total muscle volume (%TMV) |  |  |  |
| --- | --- | --- | --- | --- | --- | --- | --- | --- | --- | --- | --- | --- |
|  | Young | Old | *p* | *d* | Young | Old | *p* | *d* | Young | Old | *p* | *d* |
| All muscles | 5046.3<br>[4779.3, 5313.3] | 3999.4<br>[3732.5, 4244.4] | **<.0001** | 1.43 | 74.8<br>[72.3, 77.3] | 54.7<br>[52.2, 57.1] | **<.0001** | 2.97 | / | / |  |  |
| Anterior compartment | 193.3<br>[175.4, 212.1] | 193.1<br>[175.2, 211.9] | .9867 | 0.01 | 2.9<br>[2.7, 3.1] | 2.7<br>[2.5, 2.9] | .1380 | 0.44 | 3.9<br>[3.6, 4.2] | 4.9<br>[4.6, 5.2] | **<.0001** | -1.31 |
| Lateral compartment | 88.5<br>[76.5, 101.4] | 74.1<br>[63.1, 85.9] | .0921 | 0.56 | 1.3<br>[1.2, 1.5] | 1.0<br>[0.9, 1.2] | **.0033** | 0.89 | 1.8<br>[1.6, 2.0] | 1.9<br>[1.7, 2.1] | .3836 | -0.23 |
| Deep compartment | 177.5<br>[160.3, 195.6] | 171.7<br>[154.8, 189.5] | .6448 | 0.15 | 2.7<br>[2.5, 2.9] | 2.4<br>[2.2, 2.6] | **.0496** | 0.59 | 3.6<br>[3.3, 3.8] | 4.4<br>[4.1, 4.7] | **<.0001** | -1.08 |
| Posterior compartment | 738.5<br>[703.0, 774.8] | 654.8<br>[621.4, 689.1] | **.0010** | 1.10 | 11.1<br>[10.7, 11.5] | 9.0<br>[8.6, 9.4] | **<.0001** | 2.06 | 15.0<br>[14.5, 15.5] | 16.8<br>[16.2, 17.3] | **<.0001** | -1.20 |
| Vastus lateralis | 515.7<br>[486.2, 546.2] | 320.0<br>[296.8, 344.1] | **<.0001** | 3.36 | 7.7<br>[7.4, 8.1] | 4.4<br>[4.1, 4.7] | **<.0001** | 4.34 | 10.4<br>[9.9, 10.8] | 8.1<br>[7.7, 8.5] | **<.0001** | 2.01 |
| Vastus intermedius | 488.2<br>[459.4, 517.8] | 341.8<br>[317.8, 366.7] | **<.0001** | 2.51 | 7.3<br>[7.0, 7.7] | 4.7<br>[4.4, 5.0] | **<.0001** | 3.40 | 9.8<br>[9.4, 10.3] | 8.7<br>[8.3, 9.1] | **.0001** | 1.02 |
| Vastus medialis | 318.3<br>[295.1, 342.3] | 246.1<br>[225.8, 267.3] | **<.0001** | 1.50 | 4.8<br>[4.5, 5.1] | 3.4<br>[3.2, 3.6] | **<.0001** | 2.19 | 6.4<br>[6.1, 6.8] | 6.2<br>[5.9, 6.6] | .4668 | 0.19 |
| Rectus femoris | 212.0<br>[193.1, 231.6] | 124.8<br>[110.5, 140.0] | **<.0001** | 2.36 | 3.2<br>[3.0, 3.4] | 1.7<br>[1.5, 1.9] | **<.0001** | 3.01 | 4.3<br>[4.0, 4.5] | 3.1<br>[2.9, 3.4] | **<.0001** | 1.54 |
| Biceps femoris (short head) | 56.5<br>[47.0, 66.9] | 51.9<br>[42.8, 61.9] | .5129 | 0.22 | 0.8<br>[0.7, 1.0] | 0.7<br>[0.6, 0.8] | 0.1039 | 0.49 | 1.1<br>[1.0, 1.3] | 1.3<br>[1.2, 1.5] | .1109 | -0.41 |
| Biceps femoris (long head) | 165.9<br>[149.3, 183.3] | 127.8<br>[113.3, 143.2] | **.0010** | 1.10 | 2.5<br>[2.3, 2.7] | 1.8<br>[1.6, 1.9] | **<.0001** | 1.62 | 3.4<br>[3.1, 3.6] | 3.2<br>[3.0, 3.5] | .4746 | 0.18 |
| Semimembranosus | 176.5<br>[159.4, 194.5] | 141.4<br>[126.1, 157.5] | **.0036** | 0.97 | 2.7<br>[2.4, 2.9] | 1.9<br>[1.8, 2.1] | **<.0001** | 1.48 | 3.6<br>[3.3, 3.8] | 3.6<br>[3.3, 3.8] | .9495 | -0.02 |
| Semitendinosus | 144.7<br>[129.2, 161.0] | 98.0<br>[85.4, 111.6] | **<.0001** | 1.48 | 2.2<br>[2.0, 2.4] | 1.3<br>[1.2, 1.5] | **<.0001** | 1.97 | 2.9<br>[2.7, 3.1] | 2.5<br>[2.3, 2.7] | **.0100** | 0.67 |
| Gluteus maximus | 736.8<br>[701.4, 773.2] | 635.5<br>[602.6, 669.3] | **.0001** | 1.35 | 11.0<br>[10.6, 11.5] | 8.8<br>[8.4, 9.2] | **<.0001** | 2.29 | 14.9<br>[14.4, 15.4] | 16.2<br>[15.7, 16.7] | **.0005** | -0.90 |
| Adductor Magnus | 458.7<br>[430.8, 487.4] | 388.0<br>[362.4, 414.5] | **.0004** | 1.20 | 6.9<br>[6.5, 7.2] | 5.4<br>[5.1, 5.7] | **<.0001** | 1.93 | 9.2<br>[8.8, 9.6] | 9.9<br>[9.5, 10.3] | **.0179** | -0.61 |
| Sartorius | 85.8<br>[73.1, 97.5] | 66.0<br>[55.7, 77.2] | **.0227** | 0.76 | 1.3<br>[1.1, 1.4] | 0.9<br>[0.8, 1.0] | **.0002** | 1.12 | 1.7<br>[1.5, 1.9] | 1.7<br>[1.5, 1.8] | .6986 | 0.10 |
| Gracilis | 57.9<br>[48.3, 68.4] | 37.7<br>[30.1, 46.3] | **.0022** | 1.02 | 0.9<br>[0.7, 1.0] | 0.5<br>[0.4, 0.6] | **<.0001** | 1.35 | 1.2<br>[1.0, 1.3] | 0.9<br>[0.8, 1.1] | **.0369** | 0.54 |
| Iliopsoas | 322.6<br>[299.3, 346.7] | 235.6<br>[215.6, 256.3] | **<.0001** | 1.82 | 4.8<br>[4.5, 5.1] | 3.2<br>[3.0, 3.5] | **<.0001** | 2.53 | 6.5<br>[6.2, 6.8] | 6.0<br>[5.6, 6.3] | **.0267** | 0.57 |

Data presented as estimated marginal means [95% confidence interval] from statistical model. Cohen's *d* effect size values were estimated by dividing the differences in estimated marginal means by the residual standard deviation from the statistical model. P-values and Cohen's *d* values represent pairwise comparisons between age groups for each muscle.

In addition to the effect of age group (absolute: χ^2^(1) = 34.2, p < 0.0001; relative: χ^2^(1) = 129.3, p < 0.0001), there was a main effect of muscle (absolute: χ^2^(16) = 17090.0, p < 0.0001; relative: χ^2^(16) = 20542.3, p < 0.0001) and an age group × muscle interaction effect (absolute: χ^2^(16) = 186.5, p < 0.0001; relative: χ^2^(16) = 271.4, p < 0.0001; Table 1). Across the different muscles there was a pronounced range in the effects of age from no differences in volume of the anterior compartment of the shank in young and older adults (absolute *d* = 0.01, -0.1%, p = 0.9867; relative *d* = 0.44, -6.9%, p = 0.1380) up to a very large difference for the vastus lateralis (absolute *d* = 3.36, - 37.9%, p < 0.0001; relative d *=* 4.34, -42.9%, p < 0.0001). Older adults had significantly lower absolute muscle volume in 13 out of 17 muscles/compartments (effect sizes for significant comparisons: *d* = 0.76-3.36), indicating substantial heterogeneity across muscles (Table 1, Figure 2A). Muscle × age interaction contrasts were significant for 39 of the 136 muscle pairs that were compared (all d ≥ 1.32); specifically, there were significantly greater age-related differences in three of the quadriceps muscles compared to many of the other 16 muscles/compartments (vastus lateralis 14, vastus intermedius 10, and rectus femoris 7; Figure 3A). Conversely, the anterior compartment, deep posterior compartment, and biceps femoris short head exhibited smaller age-related differences compared to 7, 6, and 4 of the other 16 muscles/compartments, respectively (Figure 3A).

**Figure 2.**
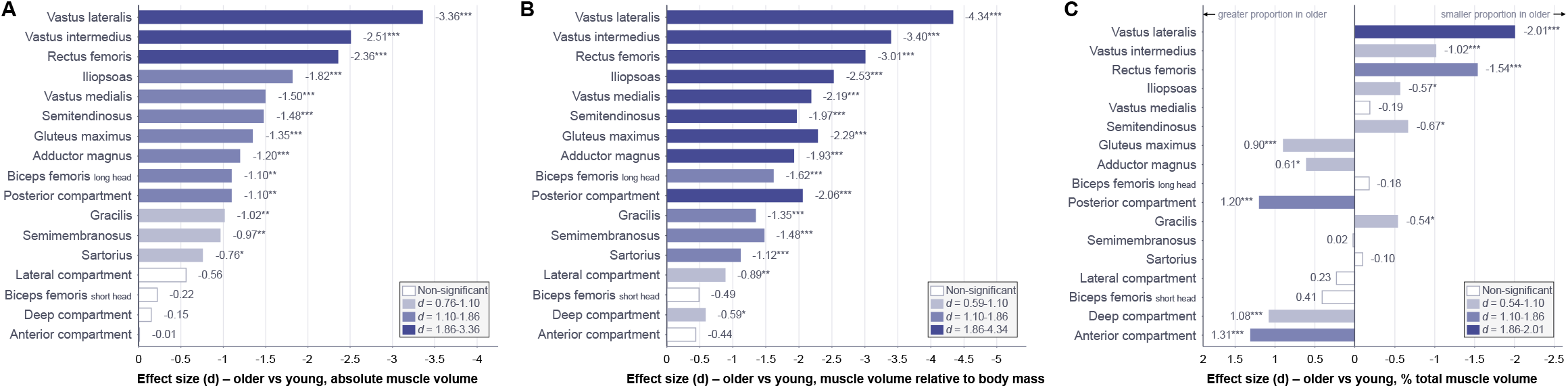
Age-related differences in the volume of 17 individual lower limb muscles and compartments, expressed as standardised effect sizes (Cohen’s d) in young and older adults. A: Absolute muscle volume (cm^3^); B: Muscle volume normalised to body mass (cm^3^·kg^−1^); C) Muscle volume expressed as a percentage of total lower limb muscle volume. Effect sizes were derived by dividing the differences in estimated marginal means by the residual standard deviation from the linear mixed effects model. Negative values denote a lower value in older adults. Muscles are ordered by the magnitude of the age-related difference in absolute volume (A). Bar shading denotes the magnitude of the effect using thresholds of |d| = 1.10 and 1.86, common to all three panels, whereas the outer values in each legend give the smallest and largest significant effect size observed in the respective panel. Open bars denote differences that were not statistically significant. *p < 0.05, **p < 0.01, ***p < 0.001 for pairwise comparisons between groups.

**Figure 3.**
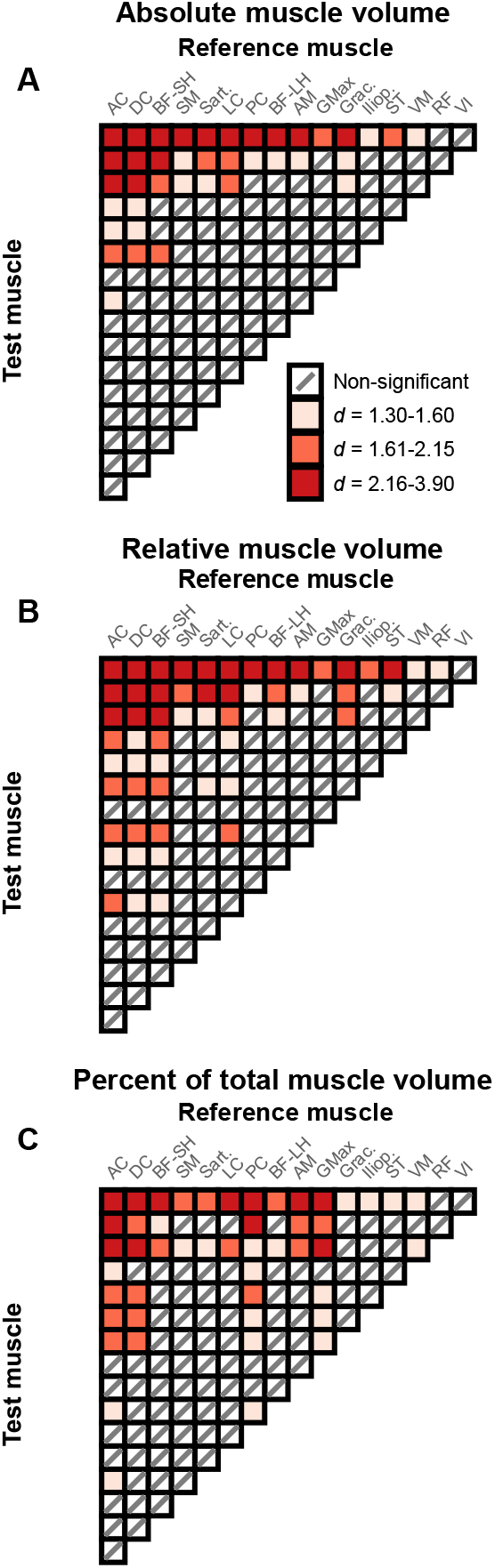
The differential effect of age on muscle volumes – interaction contrasts for absolute muscle volume (A), muscle volume relative to body mass (B), and percent of total muscle volume (C). All coloured squares represent a significantly greater effect of age in the test muscle than the reference muscle (interaction contrast, p < 0.05), whereas white squares with a diagonal line indicate no significant difference. The estimated Cohen’s *d* values for all significant effects were pooled and split into tertiles to appreciate the effect sizes of all interaction contrasts: light red, *d* = 1.30-1.60; medium red, *d* = 1.61-2.15; and dark red, *d* = 2.16-3.90. Note that all significant effects are considered large (*d* ≥ 1.30). VL = vastus lateralis, VI = vastus intermedius, RF = rectus femoris, VM = vastus medialis, ST = semitendinosus, Iliop. = iliopsoas, Grac. = gracilis, GMax = gluteus maximus, AM = adductor magnus, BF-LH = biceps femoris long head, PC = posterior compartment, LC = lateral compartment, Sart. = sartorius, SM = semimembranosus, BF-SH = biceps femoris short head, DC = deep compartment, AC = anterior compartment.

Due to older adults’ greater body mass compared to young, a lower relative muscle volume was noted in 15 out of 17 muscles/compartments with effect sizes for significant comparisons ranging from *d* = 0.59 to *d* = 4.34 (Table 1; Figure 2B). Interaction contrasts confirmed that age-related differences in muscle volume appear to be exacerbated when accounting for differences in body mass, with 56 of the 136 interaction contrasts showing significant interactions, compared with 39 for absolute volume. Specifically, greater age-related differences in relative muscle volume were noted for vastus lateralis, vastus intermedius and rectus femoris compared to 15, 11 and 8 other muscles, respectively, whilst the anterior compartment, deep posterior compartment and biceps femoris short head each showed significantly smaller age-related differences than 9 other muscles (Figure 3B).

When considering the proportion of each muscle/compartment’s contribution to total muscle volume, no significant effect of age group (χ^2^(1) = 1.0, p = 0.3124), but a significant effect of muscle (χ^2^(16) = 22876.3, p < 0.0001), and an age group × muscle interaction (χ^2^(16) = 213.5, p < 0.0001) were noted. These findings indicate that distal muscles, such as the anterior, deep, and posterior compartments of the shank, that atrophy slowly, were a greater proportion of total muscle volume in older adults compared to young, whereas proximal muscles such as the vastus lateralis, vastus intermedius and rectus femoris, that atrophy quickly were a smaller proportion of older adults’ musculature (Table 1; Figure 2C). Interaction contrasts revealed significant differences in 48 out of 136 pairwise age × muscle comparisons, with the vastus lateralis and rectus femoris contributing a smaller proportion of total muscle volume with age compared to 14 and 11 other muscles/compartments, respectively, whereas the anterior compartment, superficial posterior compartment, deep posterior compartment and gluteus maximus each contributed a significantly greater proportion compared to 6 - 9 other muscles (Figure 3C).

## Discussion

The present study compared the MRI-derived muscle volumes of seventeen lower body muscles/compartments in young and older adults. Older adults exhibited substantially lower total muscle volume, with the effect of age differing substantially between muscles. Age-related differences in absolute and relative volume were the greatest in three of the four individual quadriceps muscles (vastus lateralis: absolute *d* = 3.36, -37.9%, relative *d* = 4.34, - 42.9%; vastus intermedius: absolute *d* = 2.51, -30.0%, relative *d* = 3.40, -35.6%; rectus femoris: absolute *d* = 2.36, -41.1%, relative *d* = 3.01, -46.9%), whereas the lower leg muscles and biceps femoris short head showed smaller or non-significant differences (anterior compartment: absolute; *d* = 0.01, -0.1%, relative *d* = 0.44, -6.9%; deep compartment: absolute *d* = 0.15, -3.3%, relative *d* = 0.59, -11.1%; lateral compartment: absolute *d* = 0.56, -16.3%, relative *d* = 0.89, -23.1%, biceps femoris short head: absolute *d* = 0.22, -8.1%, relative *d* = 0.49, -12.5%). Overall, our results demonstrate substantial heterogeneity across muscles in the age-related differences in muscle volume, resulting in a redistribution of muscle mass in the lower limb of older adults compared to young, with three of the four quadriceps muscles (rectus femoris, vastus lateralis, and intermedius) representing a significantly smaller proportion, and three of the four shank compartments (anterior, deep, and posterior compartments) a significantly greater proportion of total muscle volume in older adults.

Older adults had substantially smaller total muscle volume compared to young, in absolute terms (-21%) and relative to body mass (-27%). Absolute muscle volume was lower in older adults in 13 out of 17 muscles/compartments, whereas due to older adults’ greater body mass, relative muscle volume was smaller in 15 out of 17 muscles/compartments. The exceptions, i.e. the lack of between-group differences in muscle size, were typically evident in more distal muscles: biceps femoris short head and anterior compartment of the shank (both absolute and relative) and the deep and lateral compartments of the shank (both absolute only). Across individual muscles, the magnitude of the age-related differences in muscle size varied markedly with effect sizes ranging from *d* = 0.01 to 3.36 (absolute) and *d* = 0.44 to 4.34 (relative).

Critically, extensive age group × muscle interaction contrasts confirmed that these apparent differences between muscles were statistically significant. Specifically, two clusters of differences emerged in our findings. First, three of the individual quadriceps muscles (vastus lateralis, vastus intermedius, and rectus femoris) showed the greatest age-related differences in absolute and relative muscle volume, as reflected by the greatest effect sizes, and interaction contrasts demonstrating greater age-related differences compared to at least 7 other muscles/compartments. For example, vastus lateralis in particular exhibited the largest age-related difference in absolute volume of any muscle examined, significantly greater than that of all but two of the other 16 muscles/compartments. Second, in contrast, the anterior and deep posterior compartments of the shank and the biceps femoris short head showed significantly smaller age effects than a number of other muscles. Whilst there has been relatively little attention to the age-related atrophy of individual muscles/compartments, it has previously been suggested that the quadriceps muscles undergo the greatest age-related atrophy. However, this has largely been inferred from descriptive comparisons of percentage differences within one sample [13], or between study comparisons [2], or from proportional contributions of quadriceps volume to total thigh muscle volume between young and older adults [7]. Hence, we are aware of no statistical evaluation of the of age-related differences between the lower limb muscles. Consequently, the present findings provide novel, rigorous evidence for the prior suggestion that the effect of age on individual quadriceps muscle atrophy exceeds that of many other lower limb muscles. In addition to the heterogeneity between muscles, age-related differences in muscle volume also varied between synergists acting on the same joint. Specifically, within the individual quadriceps muscles, the vastus lateralis exhibited a greater age-related atrophy in both absolute and relative muscle volume compared to the vastus medialis. Whilst two smaller studies reported no differences in the extent of muscle atrophy between individual quadriceps [7,14], the present finding that the vastus medialis exhibited less age-related atrophy is broadly consistent with Hogrel et al. [13] who also reported the smallest percentage age-related difference in vastus medialis in a large sample (n = 72). However, Hogrel et al. [13] qualitatively reported the largest percentage difference with age occurred for the rectus femoris, whereas in the present study the effect sizes and highest number of age × muscle interaction contrasts were most pronounced in the vastus lateralis. It is important to note that this might be a function of the metrics used to estimate the differences in muscle volume for each muscle. Specifically, whilst percentage differences are derived from group means alone thus not reflecting variability, standardised effect sizes express the between-group difference relative to the residual variability of the model, and scale with the absolute rather than proportional difference.

Within the hamstrings, age effects were noted in the semimembranosus, semitendinosus, and biceps femoris long head but not the biceps femoris short head, a pattern consistent with Hogrel et al. [13]. Although no age group × muscle differences were noted between individual hamstring muscles in absolute muscle volume, the relative volume of the semitendinosus showed a greater reduction with age than the biceps femoris short head. At present, the mechanisms underpinning the heterogeneity of age-related atrophy across muscles remains unclear. Interestingly, prolonged bed rest in young males also results in differential atrophy within the lower limb. Specifically, within the quadriceps, interaction contrasts demonstrated that the rectus femoris appears relatively more resistant to atrophy following unloading compared to the vasti muscles [15]. The extent to which reductions in physical activity through the lifespan contribute to the muscle-specific patterns of atrophy identified in the present study remains unresolved.

Due to the heterogeneity of age-related muscle atrophy, the distribution of muscle mass across the lower limb was significantly different between young and older adults. Critically, the proportional changes occurred in both directions, with muscles at either end of the spectrum of age-related differences showing significant interactions. Specifically, the contribution of three of the four quadriceps (vastus lateralis, vastus intermedius, and rectus femoris), semitendinosus, gracilis and iliopsoas to total muscle volume was significantly smaller in older adults. Whereas that of the two large hip extensor muscles (gluteus maximus and adductor magnus) and muscles of the shank (anterior, posterior deep, and posterior superficial compartments) were of significantly greater proportional size in older adults. The observation that some muscles were proportionally greater, and others smaller, in older compared to young adults, with interaction contrasts confirming these effects, provides strong evidence of a redistribution of muscle mass. This redistribution may have important implications for physical function and mobility in older adults. With age there appears to be a redistribution of joint torques and powers during gait, with older adults doing more work at the hip but less work at the knee and ankle compared to young individuals [16]. Additionally, sarcopenic individuals show altered muscle activation patterns during physical function tests, showing greater activation of proximal muscles and lower activation of distal muscles of the lower limb than non-sarcopenic older adults [17]. Overall, it remains unclear whether the age-related redistribution of muscle mass contributes to these functional alterations or is itself a consequence of age-related changes in movement patterns and muscle activation. Nevertheless, the muscle-specific pattern of age-related differences observed in the present study highlights the importance of considering individual muscles/compartments when designing exercise interventions to preserve physical function with ageing since loading the lower limb generically may not adequately target the most affected muscles. Given the greatest age-related differences in muscle volume were within three of the four quadriceps muscles, prudent interventions might specifically load the knee extensors, whereas the plantar flexors and dorsiflexors might require less emphasis.

### Limitations

The present study has several strengths, including the use of MRI with manual muscle segmentation, which is considered the most robust methodology for the measurement of muscle volume, a relatively large, sex-balanced cohort with a consistent level of physical activity across age groups, as well as formal testing of age × muscle interaction effects, which provides direct statistical evidence that the effect of age differs between muscles. However, some limitations must be acknowledged. First, the cross-sectional study design at a single time point means it cannot be determined whether observed differences are solely due to the effect of age or other pre-existing differences. Longitudinal data throughout the lifespan are therefore required to understand the trajectory of muscle-specific atrophy. For example, despite the vastus lateralis showing significantly greater age-related differences than most other lower limb muscles in the present study (24 vs 73 years), the hamstrings showed significantly greater atrophy than the quadriceps in a follow up of older adults from age 73 to 78 years [3]. Second, muscle volume as measured here comprises both contractile and non-contractile tissue. Intramuscular fat infiltration has been shown to increase with age such that a given muscle volume of an older individual typically contains a smaller proportion of contractile tissue compared to a young individual. Therefore, it is likely that the present results underestimate the extent of age-related differences in contractile tissue. Furthermore, although less well established, previous work has also indicated that fat infiltration with ageing may differ between muscle groups, with potentially greater fat fraction reported in the paraspinal muscles than in the leg muscles [18]. Thus, the present results might also underestimate the heterogeneity of contractile tissue differences between young and older individuals, which requires further study. Lastly, the present study did not include muscle- or joint-specific measures of function, thus preventing examination of functional consequences of age-related differences in muscle volume and the observed redistribution of muscle mass. However, in the same cohort, we have previously reported that the age-related difference in maximal isometric strength was greater in the knee extensors than the dorsiflexors [19], consistent with the muscle-specific age-related difference in muscle volume reported here. Nevertheless, systematic assessment of strength and power across a wider range of joint actions concurrently with the assessment of muscle volume is required to establish whether the magnitude of muscle-specific age-related differences predicts the magnitude of joint-specific functional differences in future work.

## Conclusions

In summary, there was a large degree of heterogeneity in the effect of age on lower limb muscle volumes. The effect of age differed significantly between muscles, with three of the four individual quadriceps muscles (and the vastus lateralis in particular), showing greater age-related differences than most other lower limb muscles, whereas the anterior and deep posterior compartments of the shank and biceps femoris short head were comparatively less affected by age. This muscle-specific atrophy with ageing results in a redistribution of muscle mass towards those muscles that show the least atrophy and away from those that show the most atrophy, which may have important functional implications and inform muscle-specific intervention strategies (e.g. greater targeting of knee extensors in resistance exercise) to maintain muscle mass and physical function with ageing.

## Acknowledgements

J.Š. was supported by Versus Arthritis Foundation Fellowship (reference: 22569).

